# Inhibiting actin polymerization does not prevent the fast block to polyspermy in the African clawed frog, *Xenopus laevis*

**DOI:** 10.1101/2020.06.18.153650

**Authors:** Maiwase Tembo, Monica L. Sauer, Bennett W. Wisner, David O. Beleny, Marc A. Napolitano, Anne E. Carlson

**Affiliations:** Department of Biological Sciences, University of Pittsburgh, Pittsburgh PA 15217

## Abstract

Fertilization of an egg by more than one sperm presents one of the earliest and most prevalent obstacles to successful reproduction. As such, eggs employ multiple mechanisms to prevent sperm entry into the nascent zygote. The fast block to polyspermy is a depolarization of the egg membrane initiated by sperm entry and is employed by diverse external fertilizers including frogs and sea urchins. For some external fertilizers, sperm entry is associated with actin polymerization during the initiation of the fast block. We therefore sought to determine whether the fast block to polyspermy in the African clawed frog, *Xenopus laevis*, requires actin polymerization. Although actin polymerization is required for sperm entry into eggs from diverse external fertilizers, including sea urchins and zebrafish, here we demonstrate that actin polymerization is not required for the fast block to polyspermy in *X. laevis*.

## Introduction

The polyspermic fertilization of monospermic eggs is embryonic lethal [1] and must be prevented. Eggs of externally fertilizing animals employ a mechanism known as the *fast block to polyspermy* to prevent fertilization by more than one sperm [1]. Within seconds of fertilization, the fast block activates a depolarization of the egg membrane [2]. Sperm can bind to, but not enter, depolarized eggs [3]. In addition to becoming depolarized, the egg also extends polymers of actin to bring in sperm at fertilization in externally fertilizing animals such as sea urchins, starfish, and zebrafish [4]. Because actin plays a prominent role bringing sperm into the of eggs of many externally fertilizing animals, we hypothesized that actin polymerization is needed for the fast block to polyspermy in the African clawed frog, *Xenopus laevis*.

## Results and Discussion

Actin polymerization is inhibited by the application of the potent drug cytochalasin B [5]. In sea urchin eggs, the actin filaments disappeared after incubation in cytochalasin B for at least 5 minutes [6]. We sought to evaluate if at similar concentrations, the inhibition of actin polymerization would prevent the fast block to polyspermy in *X. laevis* eggs. To address our question, we conducted whole cell recordings during fertilization of *X. laevis* eggs in the presence or absence of the potent actin polymerization inhibitor, cytochalasin B. Under control conditions or in the absence of cytochalasin B, we observed normal depolarizations with fertilization (Figure 1a) [2]. Surprisingly, *X. laevis* eggs inseminated in 10 µg/ml cytochalasin B also depolarized (Figure 1b). This concentration of cytochalasin B was chosen because it completely inhibited sperm entry into eggs of another external fertilizer, zebrafish [7]. Importantly, sperm entry is facilitated by the depolarization of the membrane in the external fertilizer, the sea urchin [8]. The depolarization of the sea urchin egg membrane is required for both sperm entry and the fast block to polyspermy [8].Our observation that *X. laevis* eggs depolarized despite the inhibition of actin polymerization suggested that although *X. laevis* is an external fertilizer like sea urchins, the processes involved in sperm entry and depolarization are different.

**Figure 1:**
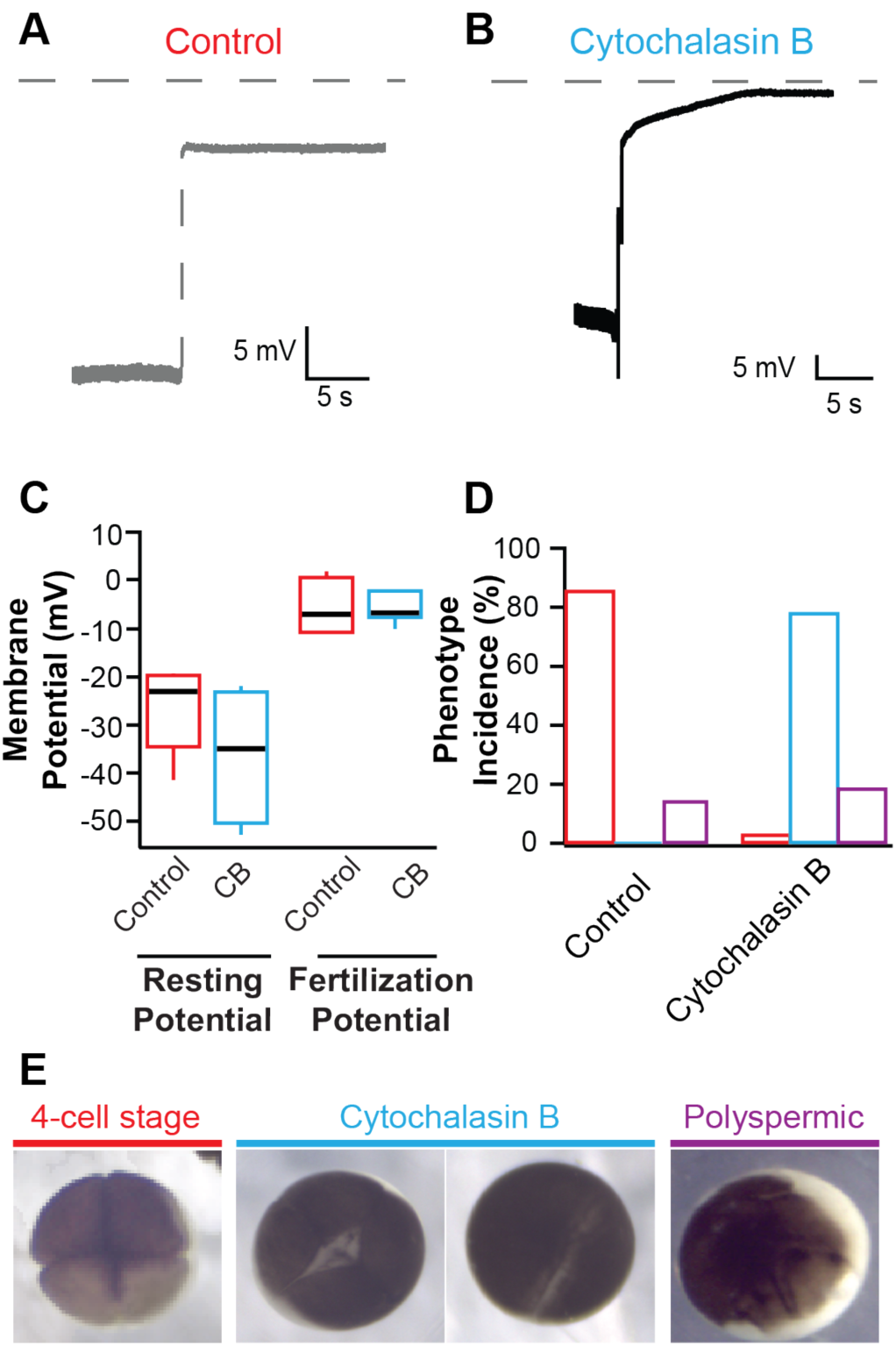
Representative whole-cell recordings made during fertilization in control conditions (a) or 10 µg/ml cytochalasin B (b). Grey dashed lines denote 0 mV. Tukey box plot distributions of the resting and fertilization potentials (c) and time to depolarization (d) in control conditions and in 10 µg/ml cytochalasin B (CB). (e) Representative images of embryos with incomplete cytokinesis, a 4-cell embryo and a polyspermic embryo with abnormal cleavage furrows. (f) Average percentage (± s.e.m.) of embryos inseminated under control conditions (red) or with 10 µg/ml cytochalasin B (blue), that developed normally to the 4-cell stage, with incomplete cytokinesis, or with abnormal cleavage furrows (n = 14). *** denotes *P* < 0.001 as determined by Student’s t-tests with the Bonferroni correction.

To examine if Cytochalasin B changed the egg’s electrical properties in a way that would affect the depolarizations recorded, we assessed the time before depolarization and egg membrane potentials. Here, *X. laevis* eggs were treated with cytochalasin B for an average of 18.6 min before depolarizations were recorded. Cytochalasin B application did not change the resting or fertilization potentials; eggs inseminated under control conditions had an average resting potential of −26.6 ± 2.9 mV (n = 10) and depolarized to 5.1 ± 1.0 mV (n = 10) which was similar to eggs inseminated in cytochalasin B whose resting and fertilization potentials were −36.1 ± 7.2 mV (n = 4) and −5.4 ± 3.1 mV (n = 4) (Figure 1c). The time between sperm addition and the onset of the depolarization was also similar between the two conditions, 6.9 ± 2.0 min for control and 6.8 ± 2.4 min for cytochalasin B (Figure 1d). These data indicate that cytochalasin B application did not alter the fast block to polyspermy. This is unlike findings in other monospermic external fertilizers such as the Spisula, the marine bivalve mollusk, whose only polyspermy block is the fast block that is capable of being inhibited at concentrations as low as 0.5 µg/ml cytochalasin B [9].

To determine whether 10 µg/ml cytochalasin B was acting on the *X. laevis* egg actin, we evaluated cleavage furrow-development of the embryos. To form cleavage furrows, *X. laevis* eggs must undergo F-actin reorganization [10], a process requiring actin polymerization. In 14 independent trials, sperm were added to groups of 10-50 eggs in 0 or 10 µg/ml cytochalasin B. 90 minutes after sperm addition, appearance of cleavage furrows was assessed for embryos in each condition. Whereas 86.6 ± 0.7% (n = 14) of eggs inseminated under control conditions developed symmetrical cleavage furrows and reached the 4-cell stage, 78.1 ± 3.4% (n = 14) eggs inseminated in cytochalasin B displayed incomplete cytokinesis (Figure 1e and f). It is possible that this concentration of cytochalasin B may effectively inhibit the actin polymerization required for cytokinesis and yet be ineffective at inhibiting the polymerization required for fertilization-evoked depolarization. However, cytochalasin B disruption of cytokinesis supports that the compound effectively inhibited actin polymerization in the *X. laevis* eggs. Not only does the egg remain fertilizable in the presence of cytochalasin B, it is capable of initiating the fast block to polyspermy unlike sea urchins or Spisula. Because the drug acts on the microfilament system of the egg, these data could have wider implications for how we think about how we think about *X. laevis* fertilization. Together, the data presented here suggest that actin polymerization is not required for the fast block to polyspermy.

## Methods

### Animals

All animal protocols were conducted using accepted standards of humane animal care, approved by the Animal Care and Use Committee at the University of Pittsburgh. Adult *X. laevis* frogs were commercially purchased (Nasco) and housed at 18°C in 12/12 hr light and dark cycle. Ovulation was induced by injection of 1,000 IU of hCG into the dorsal lymph sac. Frogs were housed at 14°C for 14-16 hours after injection. Females typically began laying eggs 0-2 hours after their transfer to room temperature. Eggs were collected onto dry petri dishes and were used within 10 min.

Testes were collected from mature *X. laevis* males following euthanasia by a 30-min immersion in 3.6 g/L tricaine, pH 7.4. To create a sperm suspension, 1/10 of a testis was minced in MR/3. Eggs were inseminated by pipetting the sperm suspension onto of the eggs bathed in MR/3. Dissected testes were stored at 4°C in L-15 medium and used within 5 days.

### Solutions

Modified Ringer’s (MR) solution used was composed of (in mM): 100 NaCl, 1.8 KCl, 2.0 CaCl_2_, 1.0 MgCl_2_, and 5.0 HEPES, pH 7.8. The MR was filtered using a sterile, 0.2-µm polystyrene filter. For fertilization-evoked depolarizations, eggs were inseminated in MR diluted to 20% MR (MR/5) for control conditions or in MR/5 with cytochalasin B added. After fertilization-evoked depolarizations were made for fertilization experiments, embryos were set aside to develop in MR diluted to 33% MR (MR/3). For developmental assays, embryos were incubated in MR/3 for control conditions or in MR/3 with cytochalasin B added. All chemicals were purchased from Thermo Fisher Scientific.

### Electrophysiology

Fertilization-evoked depolarizations were recorded as previously described [2]. Briefly, recordings were made in the whole-cell using TEV-200A amplifiers (Dagan Co.) and digitized by Axon Digidata 1550A (Molecular Devices). The data were collected with pClamp Software (Molecular Devices) at a rate of 5 kHz. Pipettes made from borosilicate glass were 8–20 MΩ resistance and filled with 1 M KCl. Membrane potentials were quantified ∼10 s before (resting) and after (fertilization) the depolarization. Data were analyzed in Excel (Microsoft) and IGOR (Wavemetrics).

### Embryonic Development Assay

Approximately 10-50 eggs were inseminated in each experimental condition and then assessed for development based on the appearance of cleavage furrows (Figure 1e). Developmental phenotypes for each frog’s eggs were scored out of the total number of eggs fertilized and developed in each condition on each experimental day.

## ACKNOWLEDGEMENTS

We thank K.L. Wozniak, R.E. Bainbridge and B.L. Mayfield for excellent technical assistance. This study was supported by the National Institutes of Health grant R00HD69410 to A.E. Carlson.

